# Chemically Tunable FOXM1-D Sensor Revealed FOXM1 Direct Influence on Cell Cycle

**DOI:** 10.1101/2023.03.01.530713

**Authors:** Kriengkrai Phongkitkarun, Porncheera Chusorn, Maliwan Kamkaew, Eric W.-F. Lam, Chamras Promptmas, Somponnat Sampattavanich

## Abstract

Forkhead box protein M1 (FOXM1) is a proliferation-associated transcription factor contributing to the G2/M phase transition of the cell cycle. Although the upregulation of FOXM1 has been observed in different cancer types, how the regulation of FOXM1 dynamically alters during cell cycles and potentially contributes to tumorigenesis is not well understood. We showed here the development and application of a tunable FOXM1-DHFR (FOXM1-D) sensor that enables surveillance and manipulation of the FOXM1 abundance. Using trimethoprim (TMP) to stabilize the sensor, we measured the kinetics of FOXM1-D production, degradation, and cytosolic-to-nuclear translocation in the G1 and G2 cell-cycle phases. By controlling FOXM1-D stability in different synchronized cell cycle pools, we found that the G1- and S-synchronized cells finished their first cell division faster, although the G2-synchronized cells were unaffected. Our analysis of single-cell FOXM1-D dynamics revealed that the two-round dividing cells had a lower amplitude and later peak time than those arrested in the first cell division. Destabilizing FOXM1-D in the single-round dividing cells enabled these cells to re-enter the second cell division, proving that overproduction of FOXM1 causes cell cycle arrest and prevents unscheduled proliferation.

## Introduction

Forkhead Box M1 (FOXM1) is a transcription factor of the Forkhead box family, consisting of 10 exons that span approximately 25 kb on the 12p13 chromosomal band (Kalathil et al., 2020). FOXM1 plays an essential role as the transcriptional regulator of a broad spectrum of downstream target genes of various biological processes, including cell proliferation, cell cycle progression, cell differentiation, DNA damage repair, migration, tissue homeostasis, angiogenesis, and apoptosis (Kalin et al., 2011, Koo et al., 2012). *FOXM1* is also an important oncogene because its dysregulation can impact all cancer hallmarks (Kalathil et al., 2020). Aberrantly high FOXM1 mRNA and protein expression has been shown in various solid tumors and correlated with advanced-stage cancer and poor prognosis (Kim et al., 2019, Li et al., 2017, Liao et al., 2018, Song and Chu, 2018). FOXM1 has also been shown to be involved with chemotherapy resistance (Chesnokov et al., 2021), making it an interesting prognostic biomarker and a potential therapeutic target for cancer treatment (Teh, 2012).

FOXM1 functions as a cell cycle regulator, controlling the expression of genes required for both G1/S and G2/M transitions. It is also essential for mitotic entry, securing chromosomal stability (Alvarez-Fernandez and Medema, 2013). To accomplish this, both its mRNA and protein abundance alters throughout the cell cycle, rising in the S phase until G2/M and remaining elevated until the late M phase (Kelleher and O’Sullivan, 2016, Liu et al., 2021). Regardless of such overall understanding, how FOXM1 contributes to tumorigenesis is still unclear. FOXM1 has been described as one of the few elevated genes during the early stages of cancer development (Chen et al., 2017, Levanon et al., 2014, Liang et al., 2021). Overexpression of FOXM1 in normal cells, like mouse embryonic fibroblasts (MEFs), triggered premature senescence (Tan et al., 2010). Undoubtedly, FOXM1 overexpression causes dysregulated cell proliferation but its mechanistic involvement during the initial stages of cancer transformation still remains enigmatic.

We introduce here a tunable FOXM1-DHFR (FOXM1-D) sensor in non-malignant MCF10A cells. By fusing the *FOXM1* gene with the mVenus fluorescent protein and the destabilizing domain from *Escherichia coli* dihydrofolate reductase (ecDHFR), the FOXM1-D sensor enables both surveillance and manipulation of FOXM1 activity. Adding trimethoprim (TMP) can inhibit the degradation of such fusion proteins (Iwamoto et al., 2010). Using quantitative imaging and time-lapse microscopy to monitor the FOXM1-D activity in the different cell cycle phases, we explored the regulation of FOXM1-D dynamics, quantifying its production rate, degradation rate, and translocation rates in both G1 and G2 phases of the cell cycle. Taking advantage of TMP as a fast approach to control FOXM1-D stability, we examined the role of FOXM1 overexpression in different cell cycle phases. Analysis of FOXM1-D dynamics at the single-cell level confirmed the role of FOXM1 as one of the critical regulators of cell cycle checkpoints. Our study elucidated how FOXM1 influences cell cycle progression and provided mechanistic insights into its potential contribution toward tumorigenesis.

## Results

### Characterization of the chemically tunable FOXM1-D sensor

To generate the FOXM1-mVenus-DHFR construct (FOXM1-D), we genetically inserted the yellow fluorescence protein mVenus and the mutant of *Escherichia coli* dihydrofolate reductase (ecDHFR) degron (Iwamoto et al., 2010) to the C-terminus of the human genomic FOXM1B-WT region (Figure 1A). We then developed an MCF10A cell line that stably expresses the FOXM1-D sensor using retroviral transduction. Due to the heterogeneity of transfection, we needed to obtain a subpopulation of cells with more homogenous FOXM1-D expression for further analysis. In short, cells were synchronized at the G2/M phase of the cell cycle using nocodazole treatment (125 nM for 14 h) and sorted into different tighter intensity bins (Supplement Figure S1A). Cell bins with strong FOXM1-D signals were chosen for further study. To monitor the FOXM1-D signal in different cell cycle phases, we used live microscopy and quantitative image processing to quantify the FOXM1-D abundance at single cells from the Venus signal intensity in both the nuclear and cytosolic compartments (Figure 1B).

**Figure 1.**
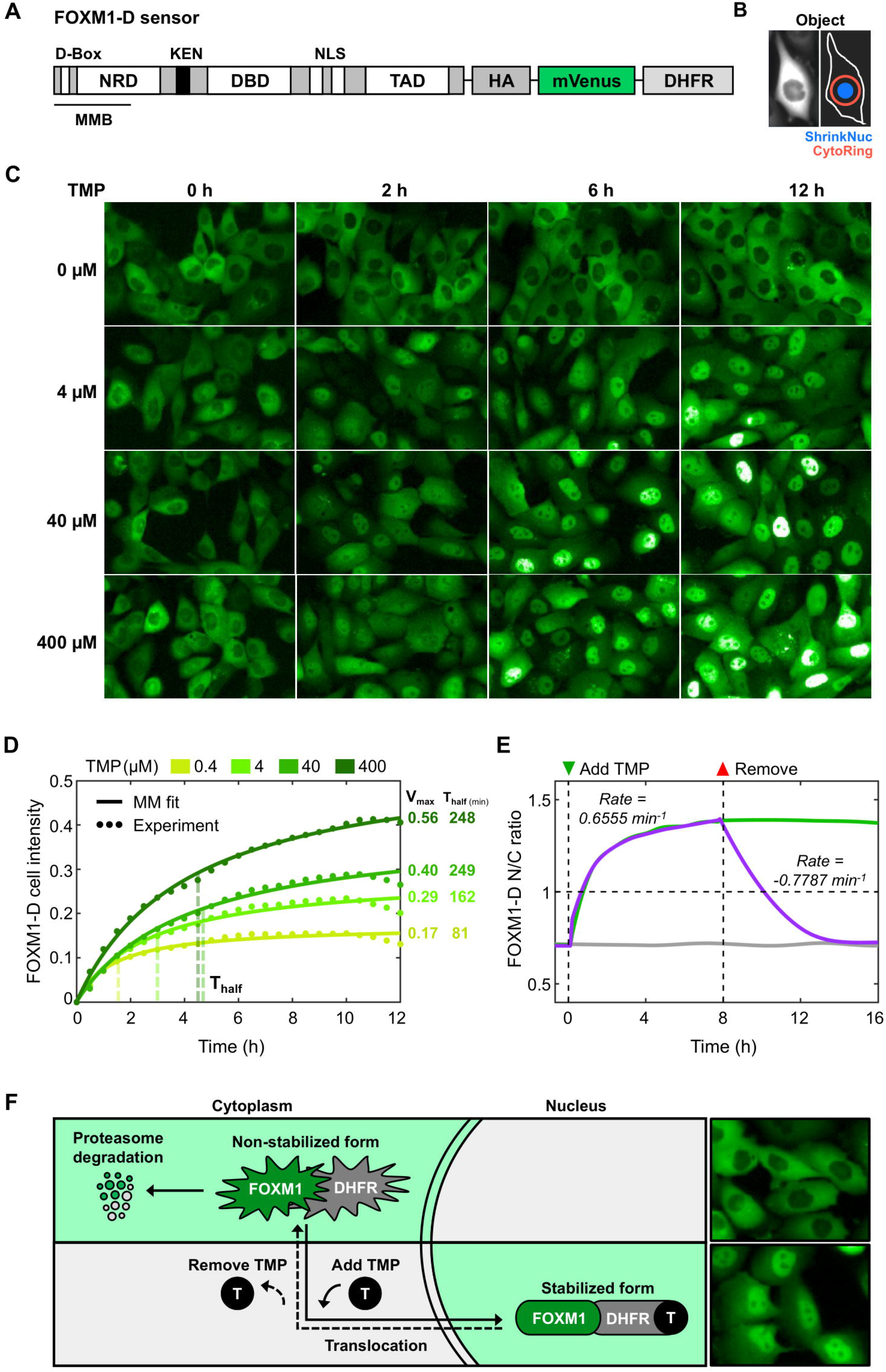
Engineering of the FOXM1-D sensor and its stability controlled by TMP. **(A)** The structure of tunable FOXM1-D sensor engineering by mVenus tagged at C-terminal. **(B)** Method used to obtain the nuclear and cytoplasmic ring regions of FOXM1-D. The nucleus was segmented by Hoechst signals and shrank for 2 pixels to obtain the nuclear region (ShrinkNuc). The cytoplasmic ring region (CytoRing) was defined by a 5 pixels-wide ring surrounding the segmented nucleus. **(C)** Images of MCF10A harboring FOXM1-D with TMP treatment (0, 0.4, 4, 40, and 400 μM) at different time points from the live-imaged assay. **(D)** Normalized FOXM1-D intensity in the whole-cell region of TMP treatment over 12 h (image every 30 min). The FOXM1-D intensities in the TMP dose-response were fitted by Michalis-Menten’s equation with 95% confidence bounds for *V_max_* and *T_half_* quantification. The mean from three independent replicates is plotted against time. **(E)** Normalized FOXM1-D activity (N/C ratio), comparing a continuous treatment of 0.4 μM TMP for 16 h (green line), with a TMP removal at 8 h (purple line). The N-to-C translocation rates were calculated and overlaid. The mean from two independent replicates is plotted against time. **(F)** The schematic concept of tunable FOXM1-D-based destabilizing system in our MCF10A model.

After obtaining a more uniform subpopulation of engineered MCF10A cells, we next characterized the tunability of FOXM1-D stability under the different protein stabilizer concentrations. In the absence of trimethoprim (TMP), we observed that the FOXM1-D sensor was mainly localized in the cytosolic compartment (Figure 1C). We hypothesized that the baseline production of FOXM1. The addition of MG-132, a protein degradation inhibitor, resulted in an elevated abundance of the FOXM1-D intensity in the cytosol (Supplementary Figure S1B), inferring that the FOXM1-D sensor must be degraded via the ubiquitin-mediated proteasome pathway (Iwamoto et al., 2010). After adding TMP, the FOXM1-D sensor rapidly (less than 3 h) translocated from the cytosol into the nucleus (Figure 1C). By quantifying the FOXM1-D intensity in the asynchronous cells, we observed that the abundance of the FOXM1-D sensor in the whole-cell compartment increases with the elevated TMP concentration (Figure 1D). The initial rate of FOXM1-D elevation in the whole cell was rapidly raised from 0-4 h, then reached steady-state sooner with the lower doses of TMP (0.4-4.0 μM). Using the Michalis-Menten (MM) modeling, we found a linear relationship between the TMP concentration and the FOXM1-D maximum level (*V_max_*) (Supplementary Figure S1C). Interestingly, the nuclear compartment was the main compartment that exhibited FOXM1-D elevation, while the cytosolic compartment showed subtle changes (Supplementary Figures S1D-E). Ultimately, this leads to elevated cytosolic-to-nuclear translocation activity with the TMP-induced FOXM1-D stabilization. Moreover, we found that the half-time (*T_half_*) of the FOXM1-D stabilization reaches a maximum level after over 4 h with TMP above 40 μM (Figure 1D). This finding implied an overly abundant supply of FOXM1-D with TMP above 40 μM, resulting in the delay of FOXM1-D cytosolic to nuclear translocation. We thus chose the 4 μM TMP concentration for our subsequent experiments to ensure the most optimal approximation of FOXM1-D kinetics.

To examine further the translocation dynamics of FOXM1-D, we monitored the translocation of the FOXM1-D sensor by calculating the N/C ratio, after TMP was added (4 μM) and later withdrawn. The FOXM1-D sensor translocated into the nucleus with the N-to-C translocation rate of 0.66 min^-1^ (Figure 1E). We next attempted to measure the kinetics of FOXM1-D re-localization back to the cytosolic compartment after TMP withdrawal. To measure this, we removed TMP (4 μM) after 8-h incubation by washing the cells with fresh media 3 times. The TMP removal caused an immediate reduction of nuclear FOXM1 with the N-to-C translocation rate of 0.7787 min^-1^ and its translocation activity vanished by over 90% after TMP was removed at 6-7 h (Figure 1E). The above results substantiate that TMP could successfully stabilize the FOXM1-D sensor, allowing the sensor to re-localize into the nuclear compartment where it functions. The simplified model of how the abundance of the FOXM1-D sensor varies and its translocation between different compartments is shown in Figure 1F.

### Examining the FOXM1-D dynamics in different cell cycle phases

Next, we were interested in investigating how the dynamics of the FOXM1-D sensor vary in the different cell cycle phases. Quantitative image-based cytometry is one of the techniques commonly used to study cell cycle at the single-cell level (Saldivar et al., 2018, Toledo et al., 2013). Based on the U-shape cell cycle plot between DNA content and EdU incorporation and the use of the Cycler computation package (Gut et al., 2015), we could classify individual cells into one of the four cell cycle segments (G1, early S (ES), late S (LS), and G2/M) and also quantify the FOXM1-D intensity in response to TMP addition (Figure 2A). First, we compared the FOXM1-D intensities to those of endogenous FOXM1 from antibody staining and of the FOXM1^WT^ sensor, FOXM1-mVenus fusion sensor without the DHFR degron (Figure 2B), all from the MCF10A cell line. Using the asynchronous MCF10A cells with a FOXM1-D sensor, we determined the change of FOXM1-D intensity in the four major cell cycle phases after 8 h of treatment with TMP. Since FOXM1 functions as a transcription factor mainly in the nuclear compartment, we focused our measurement of FOXM1-D intensity mainly in the nuclear compartment. Based on the single-cell fluorescence images of EdU and DNA-stained MCF10A cells harboring the FOXM1^WT^ sensor, we observed an initial increase of the FOXM1^WT^ sensor in the ES phase, and its peak during the LS and G2/M phases (Figure 2C). This result can also be better visualized along the continuous cell cycle trajectory plot, showing its gradual increase and maximum intensity in the G2/M phase across all conditions (Figure 2D). Specifically, the endogenous FOXM1 and FOXM1^WT^ sensors increased by over two folds in the G2/M compared to its baseline amount in G1 (Supplementary Figure S2A, blue and red lines), while the FOXM1-D sensor under TMP treatment increased by 1.6 folds (Supplementary Figure S2A, green line). Moreover, with TMP addition, the abundance of the stabilized FOXM1-D sensors increased by 1.8 folds during G1 and 3.2 folds during G2 compared to its baseline amount of non-stabilized form during G1 (Figure 2E, dark green line). The non-stabilized FOXM1-D demonstrated only a slight increase of 1.1 folds in G2 relative to its baseline level in G1 (Figure 2E, green line).

**Figure 2.**
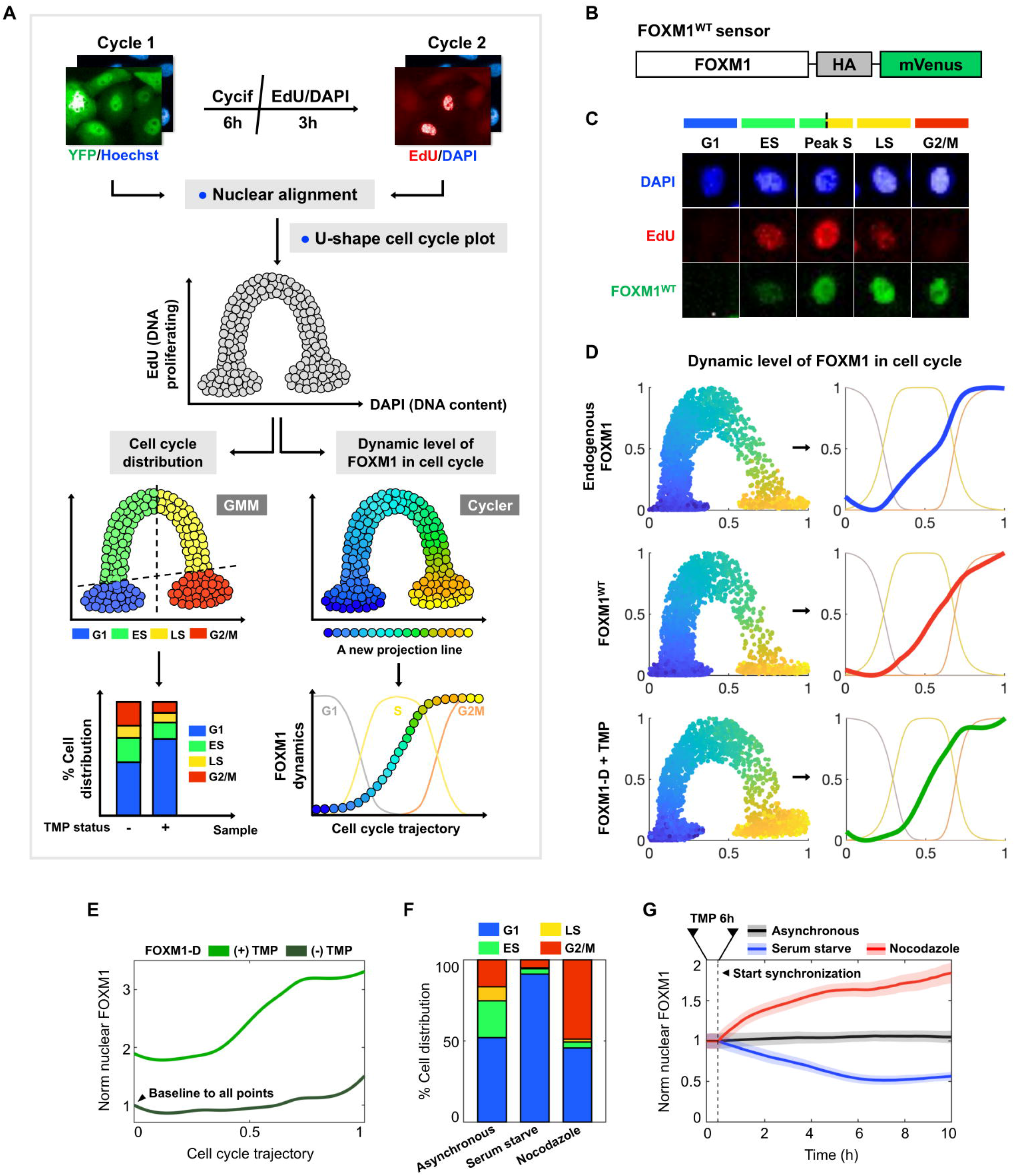
Analysis of cell cycle profiling and FOXM1 trajectory along the cell cycle. **(A)** Quantitative image-based analysis in cell cycle distribution and FOXM1 trajectories. **(B)** The FOXM1^WT^-mVenus construct **(C)** Images of the FOXM1 in cell cycle profiling in the FOXM1^WT^-mVenus MCF10A. **(D)** FOXM1 dynamics in cell cycle trajectories demonstrated in endogenous FOXM1, FOXM1^WT^, and FOXM1-D with TMP. Mean of single-cell trajectories are shown (n > 1,000 per condition). **(E)** Normalized nuclear FOXM1-D dynamics in cell cycle trajectories. The level of FOXM1-D with TMP was normalized by its baseline amount in G1 of non-stabilized forms (dark green line). The non-stabilized FOXM1-D was normalized by its baseline amount in G1 (light-green line). Mean of single-cell trajectories are shown (n > 1,000 per condition). **(F)** Cell cycle distribution of FOXM1-D overexpressing cells, after synchronization by serum-starvation and nocodazole (125 nM) for 10 h (n = 3 per condition). **(G)** Nuclear FOXM1-D dynamics changes during cell synchronization corresponding in Figure 2F after TMP addition for 6 h in live-cell imaging assay. The mean and SD from three independent replicates are plotted against time.

We next investigated the effect of drug-induced cell synchronization on the FOXM1-D dynamics. Specifically, we examined how nocodazole and serum starvation would affect FOXM1-D changes. To answer this, we first stabilized FOXM1-D by adding TMP for 6 h and synchronized the cells at the G1 phase by serum starvation and G2/M by nocodazole treatment (125 nM). The dynamical change of FOXM1-D was then monitored using live microscopy for 10 h. We confirmed using the single-cell measurement of EdU incorporation and DNA content that the serum-starved and the nocodazole-treated cells were indeed arrested in the G1 and G2/M phases (Figure 2F). We then compared the nuclear intensity of the FOXM1-D sensor between the two arrested points. As predicted, serum-starved cells exhibited a reduction of nuclear FOXM1-D intensity, while the nocodazole-treated cells showed an increase in the nuclear FOXM1-D intensity (Figure 2G). This data confirmed that the abundance of FOXM1-D sensor changes with cell cycle states similar to what we expect from the endogenous FOXM1. Moreover, we were able to tune the expression of the FOXM1-D sensor in the different phases of the cell cycle chemically using TMP.

### FOXM1-D sensor enables the quantification of FOXM1 kinetics during different cell-cycle phases

FOXM1 has been known to be cell cycle-dependent, although its degradation and cytosolic-to-nuclear transport regulation has not been thoroughly understood. With the conventional bulk assay, the measurement of FOXM1 transport kinetics in the different cell-cycle phases would not be possible due to the asynchronous nature at the single-cell level. Even with cell synchronization, the single-cell heterogeneity of protein expression would also complicate the measurement of FOXM1 kinetic parameters. Taking advantage of our chemically tunable FOXM1-D sensor, we sought to examine how FOXM1 stability varies during the different cell-cycle phases. While FOXM1 is usually undetectable in the G1 phase of the cell cycle, the FOXM1-D sensor can be detectable both in the cytosolic and nuclear compartments throughout different cell-cycle phases, offering the possibility to measure its degradation and translocation dynamics (Figure 3A). We modeled the kinetics of the FOXM1 transport between the different cellular compartments using the two-compartment transport model (Figure 3B), as described in the core model (Equations 1 and 2). In brief, the synthesis of a new FOXM1-D molecule is described by the rate parameter *k_s_*. The degradation of FOXM1-D in the cytoplasm in the presence of TMP is explained by the rate parameter *k_d_*. The transport of stabilized FOXM1-D molecule into the nucleus is defined by the rate parameter *k_cn_*. Relocation of stabilized FOXM1-D back into the cytoplasm is defined by the rate parameter *k_nc_*. We examined these rate parameters in the synchronized cells at either G1 or G2 cell-cycle phases. To determine the value of *k_s_, k_d_, k_cn_*, and *k_nc_* in either G1 and G2 phases, we estimated the relative ratio (*R*) of the two cell cycle phases (G2 over G1) for each rate parameter and later used these relative values as the constraints in our modeling process. To measure the rate of FOXM1-D synthesis *(k_s_)* (Equation 3), we treated the cells with MG-132, a protein degradation inhibitor. The FOXM1-D intensity was then measured in both compartments without TMP addition. The FOXM1-D synthesis rates in the G1 and G2 phases were found to be 0.0018 unit/min and 0.0128 unit/min, respectively, giving the FOXM1-D synthesis rate ratio (*R_s_*) of 7.11-fold higher in G2 over G1 (Figures 3C and 3E, upper panel). We then measured the degradation rate (*k_d_*) (Equation 4) by first stabilizing the FOXM1-D sensor using TMP for 4 h and then inhibiting the protein production using Cycloheximide (CHX) treatment. We assumed that the FOXM1-D degradation occurred only in the cytosol and that the FOXM1-D translocation was also affected by this process. We thus estimated the cytosolic FOXM1-D abundance in the presence of TMP. The degradation rate of the FOXM1-D sensor during the G1 and G2 phases was found to be 0.0004/min and 0.0031/min, respectively. This result gives the ratio of FOXM1-D degradation rate (*R_d_*) of 7.75-fold higher in the G2 phase than in the G1 phase (Figures 3D and 3E, middle).

**Figure 3.**
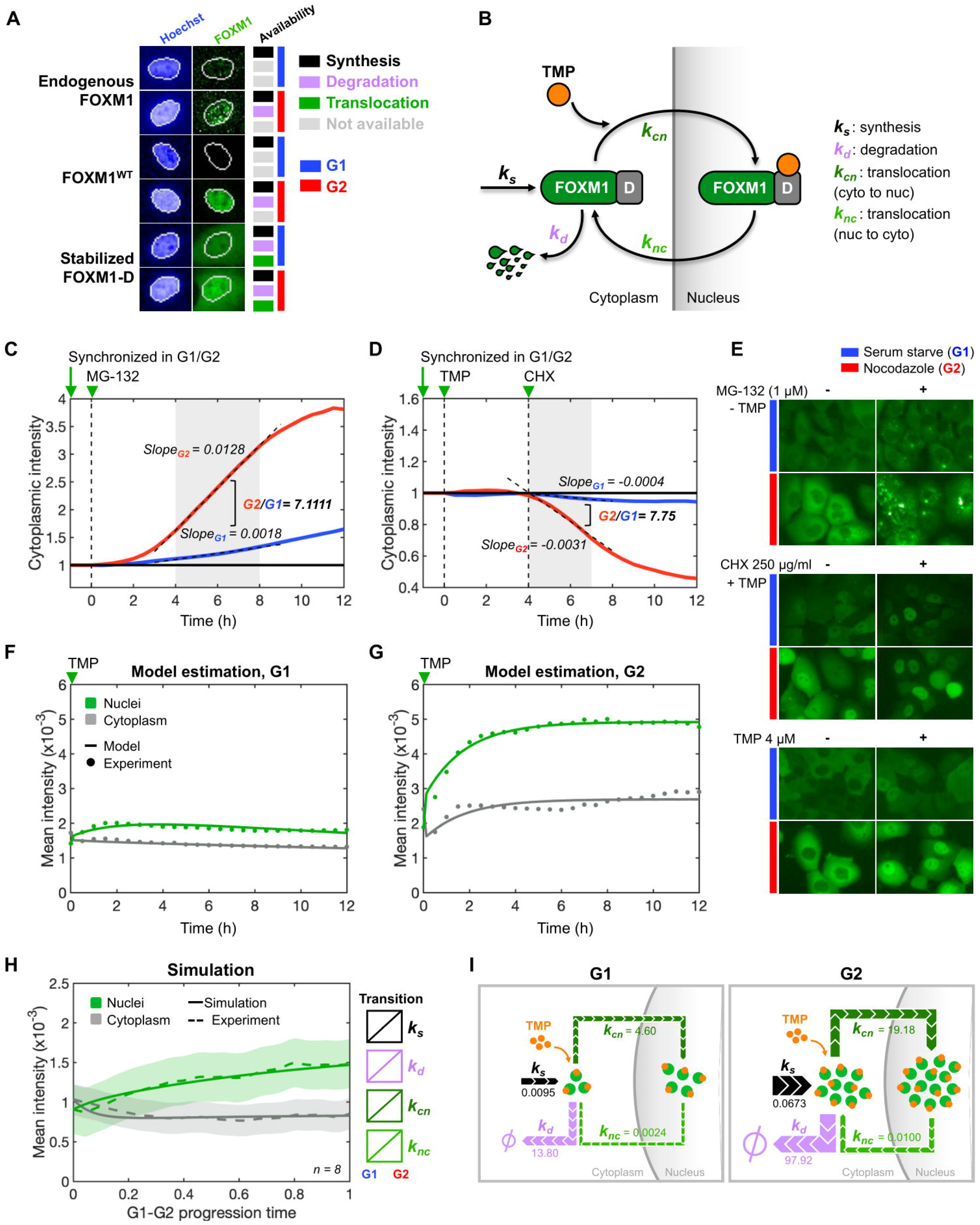
Dynamic modeling of the FOXM1-D in the G1 and G2. **(A)** Possibility and availability to use the endogenous FOXM1, FOXM1^WT^, and FOXM1-D as a tool for measuring its production, degradation, and translocation dynamics in the cell cycle. **(B)** Schematic of FOXM1-D two-compartment transport model between the different cellular compartments. **(C-D)** Normalized cytoplasmic FOXM1 intensity in **(C)** MG132 treatment for 12 h and **(D)** Cycloheximide (CHX) treatment for 8 h. Cells were synchronized in either G1 using serum starvation (blue line) or G2 using nocodazole treatment (red line) before being treated. Then, the synthesis or degradation rates in each condition were calculated by the slope between 4-8 h after the treatment. The mean from two independent replicates is plotted against time. **(E)** Images of MCF10A harboring FOXM1-D in various treatment conditions from the live-imaged assay. **(F-G)** Model estimation of FOXM1-D compartment dynamics in **(F)** G1-synchronized and **(G)** G2-synchronized conditions, compared with the experimental observation after treatment with TMP. The mean from two independent replicates is plotted against time. **(H)** Simulation of FOXM1-D compartment dynamics in the linear-rate transitions between G1 and G2 phases, compared with the experimental data in live-single cell imaging (n = 8 cells). **(I)** FOXM1-D kinetics rates in the cells between the G1 and G2 phases of the cell cycle.

To estimate the FOXM1-D translocation rates (*k_cn_, k_nc_*), we used the rate ratios *R_s_* and *R_d_* from the previous experiments as the constraints to solve Equations 5 to 8, with ±10% restricted boundaries. The FOXM1-D intensity was measured after TMP addition to enhance the translocation process. The best-fit trajectories during the G1 and G2 phases are shown in Figures 3F-G, with the corresponding images shown in Figures 3E, lower panel. We found the FOXM1-D translocation rate ratios from nuclear to the cytosol and vice versa (*R_cn_*, *R_nc_*) to be approximately 4-fold higher in G2 than in G1. Table 1 summarizes the estimation of relative rate ratios for all parameters.

**Table 1.** FOXM1-D model estimation of relative rate ratios (G2/G1)

|  |  |  |  |
| --- | --- | --- | --- |
| $R_s$ | $R_d$ | $R_{cn}$ | $R_{nc}$ |
| 7.1111 | 7.7500 | 4.1701 | 4.1599 |

We next performed a model simulation to validate the estimated parametric values (*k_s_, k_d_, k_cn_, k_nc_* in G1 and G2 phases) in three conditions: (1) when we set the relative rate ratios using results from table 1 as the modeling constraints with a boundary limit of 10% (Equations 1 and 2), (2) when we measured the FOXM1-D intensity from our single-cell live-imaging dataset as it progressed from G1 to G2 and (3) when we attempted to model the FOXM1 kinetics throughput cell cycle, by assuming a linear transition for all kinetic parameters from G1 to G2. Interestingly, we found that our assumption to have linear rate transitions between the G1 and G2 phases of the cell cycle was able to deliver simulation results that match closely with our experimental data (Figure 3H). We observed a dynamical increase of the nuclear FOXM1-D during the G1-to-G2 transition, while the cytosolic FOXM1-D decreased during this time window. After we extracted the parametric value of *k_s_, k_d_, k_cn_*, and *k_nc_* from the model simulation, we discovered that the FOXM1-D synthesis, degradation, and translocation rates were generally higher in G2 than in the G1 phase, as summarized in table 2. Furthermore, the FOXM1-D nuclear import rates (*k_cn_*) in G2 and G1 were 19.180 min^-1^ and 4.599 min^-1^, respectively, substantiating its roles as an essential transcriptional factor during the G2 phase. We summarized the kinetic parameters of the FOXM1-D dynamics between the G1 and G2 phases in Figure 3I.

**Table 2.** FOXM1-D model simulation

| Phase | $k_s(\text{U} \cdot \text{min}^{-1})$ | $k_d(\text{min}^{-1})$ | $k_{cn}(\text{min}^{-1})$ | $k_{nc}(\text{min}^{-1})$ |
| --- | --- | --- | --- | --- |
| **G1** | 0.0095 | 13.7954 | 4.5995 | 0.0024 |
| **G2** | 0.0673 | 97.9274 | 19.1802 | 0.0100 |

### Impact of FOXM1 overexpression on cell cycle and cell fate decision

Because the tunable FOXM1-D system allows conditional control of FOXM1 abundance, we were interested in examining the impact of FOXM1 overexpression during different cell-cycle phases. First, the FOXM1-D-expressing MCF10A cells were synchronized in three ways: 1) Serum-deprived G1 cells, 2) S-synchronized cells using double thymidine blockade, and 3) G2-synchronized cells using nocodazole. Cells were then released to undergo a normal cell cycle together with the addition of TMP (4 μM), aiming to stimulate FOXM1 overexpression in the different cell cycle phases. Cell-cycle characteristics and the FOXM1-D dynamics at single cells were monitored using live microscopy for 48 h (experimental process summarized in Figure 4A). Cytokinesis events (n=30) were manually tracked to annotate cell division characteristics (Figure 4B). In our 48-hour observation period, cells synchronized in G1 and S phases divided only 1-2 times, whereas G2 cells could divide up to 3 times (Figure 4B). When we quantified and compared the onset time of 1^st^ cell division, cells that started in G1 and S phases with overexpressed FOXM1-D (with TMP addition) split significantly faster than those without FOXM1-D (no TMP) for over 4 h (Figure 4C, with median comparison). Interestingly, we did not observe a similar onset time difference between cells synchronized in the G2 phase, with or without TMP. We hypothesized that the overexpression of FOXM1 in G2-synchronized cells may not rise fast enough to compete with the cell-splitting events within our 4-hour monitoring period.

**Figure 4.**
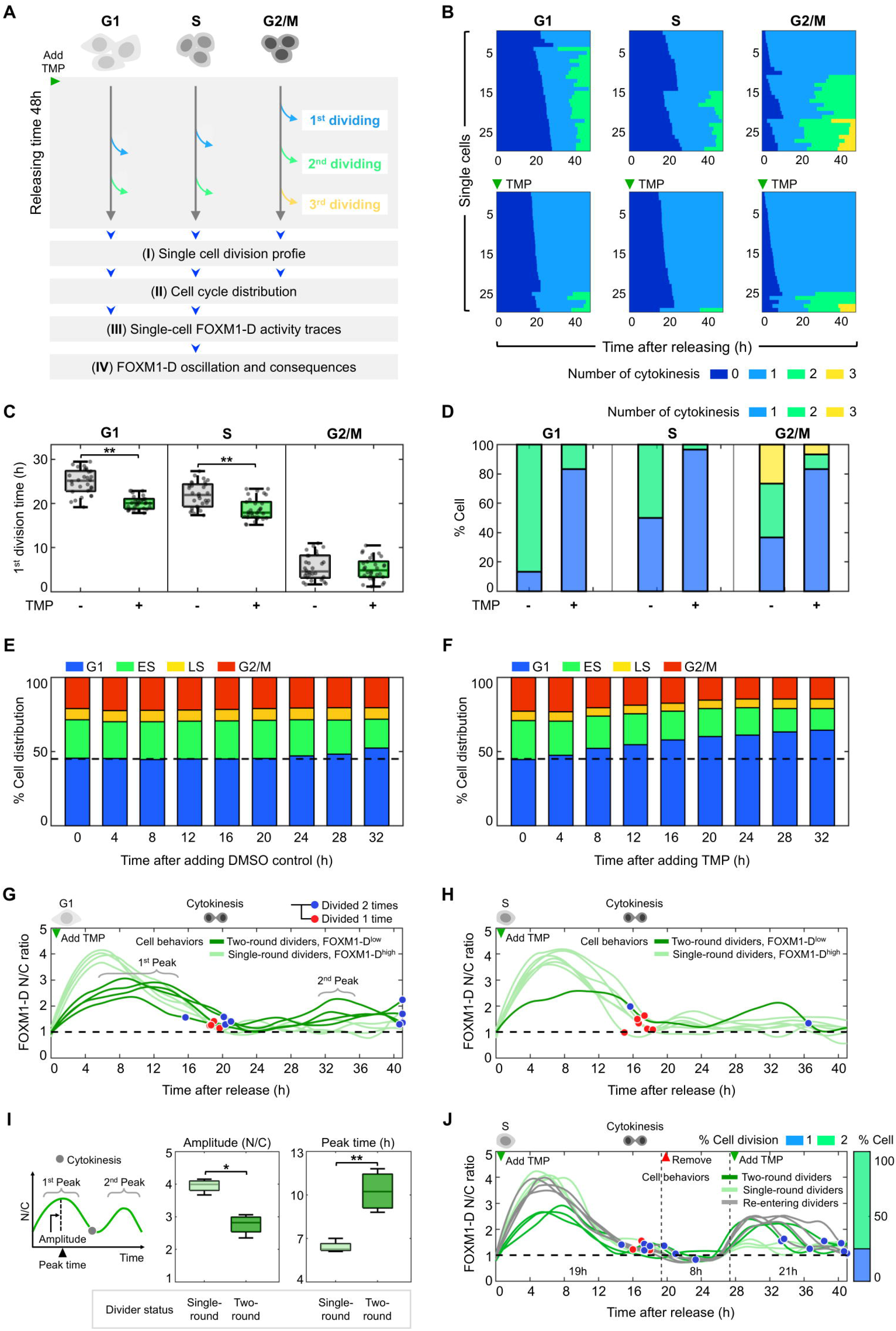
Effect of FOXM1-D overexpression on cell division and cell fate decision. **(A)** Experimental workflow to study the effect of FOXM1-D overexpression on cell division and cell fate decision **(B)** Division profile from single-cell tracking of FOXM1-D overexpressing cells in different cell cycle phases. Cells were first synchronized, then released to enter the regular cell cycle, together with the addition of 4 μM TMP to overexpress FOXM1-D. Cytokinesis events were annotated for 48 hr after releases. Each row represents the division profile of an individual cell over time. Colors change upon cytokinesis event. Cells are grouped by their total number of cytokinesis and ordered by the timing of their first cytokinesis. **(C)** Quantification and comparison of the onset time of 1^st^ cell division between FOXM1-D overexpressing cells (with TMP) and negative control (no TMP) in different phases of the cell cycle (n = 30 cells per condition). ** *P* < 0.01. **(D)** Fraction of cells with several cytokinesis events, compared between FOXM1-D overexpressing cells (with TMP) and negative control (no TMP) in different phases of the cell cycle (n = 30 cells per condition). **(E-F)** Cell cycle distribution of asynchronous FOXM1-D expressing cells after adding DMSO control **(E)** and TMP **(F)** in different time points. The mean from two independent replicates is plotted against time. **(G-H)** Single-cell traces of FOXM1-D activity (N/C ratio) in FOXM1-D overexpressing cells (with TMP) after cell cycle releasing from G1 phase **(G)** and S phase **(H)**. Traces were colored dark green for two-round dividers with low FOXM1-D activity (n = 4 cells in **(G)** and n = 1 cell in **(H)**), while single-round dividers with high FOXM1-D activity were colored in light green (n = 4 cells in **(G)** and n = 6 cells in **(H)**). The cytokinesis time events were marked with blue (divided two times) or red dots (divided one time). **(I)** Distributions of FOXM1-D first peak amplitude and peak time, compared between two-round dividing population (dark green color) and single-round dividing population (light-green color), on single-cell traces from Figure 4G. (n = 4 cells per condition). * *P* < 0.05 and ** *P* < 0.01. **(J)** Single-cell traces of FOXM1-D activity (N/C ratio) in FOXM1-D expressing cells, after cell cycle releasing from S phase. The TMP treatment schedule manipulated the FOXM1-D level to mimic the FOXM1 activity during a normal cell cycle. Traces were colored dark green for two-round dividers with low FOXM1-D activity (n = 3 cells), while single-round dividers with high FOXM1-D activity were colored in light green (n = 3 cells). Re-entering dividing cells were colored in grey (n = 4 cells).

Additionally, we observed that cells with FOXM1-D overexpression in either G1, S, or G2 phases could not proceed normally to the following cell division, defined here onwards as single-round dividers (G1: 83.3%, S: 96.7%, G2: 83.3%) (Figure 4D). Therefore, we proposed that following their initial cell division, these single-round dividers were arrested in the following G1 cell-cycle phase. This is consistent with the low abundance of nuclear content (Hoechst sum intensity) after the first cell division, as compared to those from cells that could proceed to the next cell divisions (the two-round dividers) for both G1-synchronized (Supplementary Figures 4A) as well as S-synchronized cells (Supplementary Figures 4B). This finding supported our observation from the asynchronous cells where we observed cell cycle arrest in G1 following TMP addition (Figures 4E-F). Because some cells could divide at an earlier time point, we started to see an increase in G1-arrested cells as early as 8 h with its accumulation up to 32 h following TMP addition.

We next compared the single-cell temporal dynamics of FOXM1-D between the G1 and S-synchronized populations (Figures 4G-H). Overall, we observed heterogeneity of the FOXM1-D N/C ratio over 2-fold during the first peak of FOXM1-D before the first cytokinesis event (G1 at 20 h, S at 18 h). After the first peak, the FOXM1-D N/C ratio remained low at ~1.2 to 1.5-fold. Interestingly, a few cells could enter the second cell division following another peaking of FOXM1-D activity (Figures 4G-H). Interestingly, we could separate two distinct cell subpopulations based on whether they could enter the second-round cell division. We attempted to compare the FOXM1-D N/C amplitude and the peak time prior to the first cytokinesis event (1^st^ peak amplitude in Figures 4G, 4I). The N/C pulse amplitude was 1.43-fold lower in the two-round dividers than in the single-round dividers. Moreover, the N/C peak time of two-round dividers also happened later than the single-round dividers by approximately 4 h (median comparison). Collectively, we observed that the single-round dividers exhibited taller and sooner peaks of FOXM1-D activity after TMP stabilization (designated as FOXM1-D^high^ in light-green traces, Figures 4G-H), while the two-round dividers showed less pronounced peaks of FOXM1-D activity at slightly later timepoint (designated as FOXM1-D^low^ in dark green traces, Figure 4G-H).

Finally, we then attempted to manipulate the level of FOXM1-D during cell cycle progression to confirm its influence on cell fate decision. Specifically, we attempted to restore the degradation of FOXM1-D of single-round dividers, hoping to force cells back into the following cell division (Figure 4J). Cells were initially synchronized in the S phase, then released with the addition of TMP (4 μM) for 19 h to achieve the first-round cell division time at 16-18 h post-releasing (Figure 4J). After the first division, cells in the G1 phase were washed 3 times using fresh media to remove TMP. Cells were subsequently cultured for another 8 h in complete media without TMP to allow degradation of the FOXM1-D sensor, as shown in Figure 1E. We observed that the N/C ratio initially decreased below 1 (Figure 4J), indicating the FOXM1-D sensor re-localization back to the cytosolic compartment after TMP removal. This process prevented the overexpression of FOXM1-D, preserving only the endogenous FOXM1 level during the 8-h G1 phase. Following that, TMP was used to re-stabilize the FOXM1-D sensor to induce the FOXM1-D overexpression again for the following 21 h, mainly to cover the following cell division. In the cell population that could enter the second division (two-round dividers), the N/C ratio immediately increased to 2.5 (labelled in dark green line; Figure 4J). Additionally, we observed that the original single-round dividers, defined by 1^st^-N/C amplitude and the peak time, were also able to re-enter this second cell division (labelled in grey line; Figure 4J), as indicated by the increase in N/C ratio post-TMP re-stabilization. This result suggested that while FOXM1-D overexpression in the G1 and S phases speeds up subsequent cell division, continuing the FOXM1-D overexpression through the G1 phase in the daughter cells could promote cell cycle arrest. This finding confirmed that maintaining the appropriate balance of FOXM1 synthesis and degradation is crucial for successful cell cycle progression.

## Discussion

We reported here the development and characterization of the FOXM1-D sensor enabling the monitoring and tuning of FOXM1 abundance using TMP. Together with cell synchronization, the FOXM1-D sensor allowed us to quantify the rate of changes of FOXM1 in cell cycles, and to investigate the impact of FOXM1 overexpression on cell cycle dynamics. Several studies reported the cytoplasmic localization of FOXM1 during late G1 and S phases before it was phosphorylated by cyclin E-CDK2/ Raf-MEK-ERK and translocated to the nucleus prior to the G2/M phase (Bektas et al., 2008, Kelleher and O’Sullivan, 2016). However, cytosolic endogenous FOXM1 protein is hardly measurable. The FOXM1-D sensor could be detectable in the cytosolic compartment of the MCF10A cells possibly due to (1) excessive FOXM1-D production (Kustikova et al., 2003, Wu et al., 2003), (2) limited protein degradative machinery in the cells, and (3) the preferential localization of DHFR fusion protein in the cytosol (Arkowitz et al., 1993, Kraut et al., 1986). Using TMP to tune its abundance, we were able to use the FOXM1-D biosensor to measure the FOXM1 transport kinetics between the nucleus and the cytoplasm. Finally, the FOXM1-D sensor enabled rapid regulation of FOXM1 abundance in any cell cycle phase. The addition of TMP showed the accumulation of FOXM1-D that reached a steady state within 4 h. Likewise, the removal of the TMP ligand resulted in the reduction of the FOXM1-D sensor with a half-life of ~3 h. While being able to overexpress FOXM1 at 2-3 fold over the baseline line, the FOXM1-D sensor enables faster overexpression of FOXM1 than the conventional exogenous expression technique (Hu et al., 2019).

Protein homeostasis depends on the complex interplay between protein synthesis and degradation that requires precise execution of each step to maintain its level and activity during cellular signalling and response (Alber and Suter, 2019). In our system, the stabilization of FOXM1-D by TMP resulted in a disruption of its baseline level in the cell cycle by affecting the synthesis and degradation process (Figure 2E), leading to subsequent cellular abnormalities and responses (Figures 4A-F). To explore the dynamical change towards this disruption, the kinetic rate of FOXM1-D was revealed for the first time in this study using the two-compartment transport model. Our tunable FOXM1-D system showed that under the universal promoter controlling of FOXM1-D production, the synthesis (accumulation) rate was 7.11-time (*R_s_*) higher in G2 over G1 phase (Table 1), which demonstrated the differential baseline production between these two cell-cycle phases. To maintain the FOXM1-D protein homeostasis, the newly synthesized FOXM1-D was found to be degraded at an equivalent rate ratio, 7.75-time (*R_d_*) higher in G2 over G1 (Table 1). In addition, the newly synthesized FOXM1-D preferred to be transported from cytosol to the nucleus with the significantly higher cytosolic-to-nuclear rate (*k_cn_*) in G2 phase (Table 2). Interestingly, during the G2 phase, we also observed the rapid degradation rate (*k_d_*) in the cytoplasm, demonstrating the excess amount of FOXM1-D remaining in the cytoplasm. We hypothesized that the excess FOXM1-D was possibly caused by the overproduction of FOXM1-D (4 μM TMP) and limited by some nuclear transporter protein partners such as nucleophosmin (NPM) in the MCF10A cells (Bhat et al., 2011). Further comparison with other computational models of the cell cycle (Ali Abroudi, 2015), as well as the addition of other FOXM1 regulators to the models that could affect FOXM1 synthesis, degradation, and translocation, may further improve the approximation of FOXM1 transport kinetics throughput cell cycle progression.

While it is commonly known that FOXM1 is cell cycle-dependent, how FOXM1 affects cell cycle and cell fate decisions that may lead to tumorigenesis is still not carefully investigated. By employing our tunable FOXM1-D system to simulate FOXM1 overexpression in the different cell cycle phases, we examined this question carefully. In our MCF10A model, a non-malignant breast epithelial cell line (Puleo and Polyak, 2021, Qu et al., 2015), we found that overexpressing FOXM1-D activity in the G1 and early S phase-synchronized cells accelerated cell proliferation during the first cell cycle but caused the daughter cells to arrest in G1 in the following cell division. This is consistent with the observation of cell enlargement and the accumulation of G1 subpopulation in asynchronous MCF10A cells that were incubated for the long term with TMP (Supplementary Figures 4C-D). These phenotypes mirror the initial characteristics of senescence cells (Gonzalez-Gualda et al., 2021, Neurohr et al., 2019), which was also demonstrated in normal hepatocytes of FOXM1 enriched cells (Baranski et al., 2015). Interestingly, Tan et al. also observed a similar phenomenon in early passages of transgenic (TG) mouse embryonic fibroblasts (MEFs) with elevated FOXM1 (Tan et al., 2010). This discovery demonstrated th e role of FOXM1 overexpression as a primary anti-proliferative mechanism in normal cells, causing replicative stress or senescence to stop the uncontrolled cell growth that may result in cancer formation (Braig and Schmitt, 2006, Courtois-Cox et al., 2008).

Finally, our study demonstrates that the increasing pattern of FOXM1-D on mother cells during the G1/S transition marks a bifurcation point for the daughter cell proliferation, where either starts the next cell cycle or transiently arrests (Fig 4G-J). We identified a threshold of FOXM1-D activity in the mother cells as a crucial factor for predicting the possibility of subsequent cell division. In summary, our system confirmed the oncogenic properties of the FOXM1 gene to accelerate cell proliferation by its overexpression in G1 and S-synchronized cells (Fig 5B-C). Moreover, the completion of FOXM1 degradation at the end of mitosis critically defines how daughter cells proceed in the next cell cycle phase (Fig 5C), which was reported to be mainly degraded by the APC/C and Cdh1 complex (Park et al., 2008, Zhou et al., 2016). We also demonstrated that the insufficient degradation of FOXM1-D following mitosis led to the G1 arrest in daughter cells, causing abnormal proliferation (Figure 5B). Future studies are still needed to examine the dynamics of FOXM1-D in cancer cell line models and to prove the connection of FOXM1 dysregulation towards tumorigenesis.

**Figure 5.**
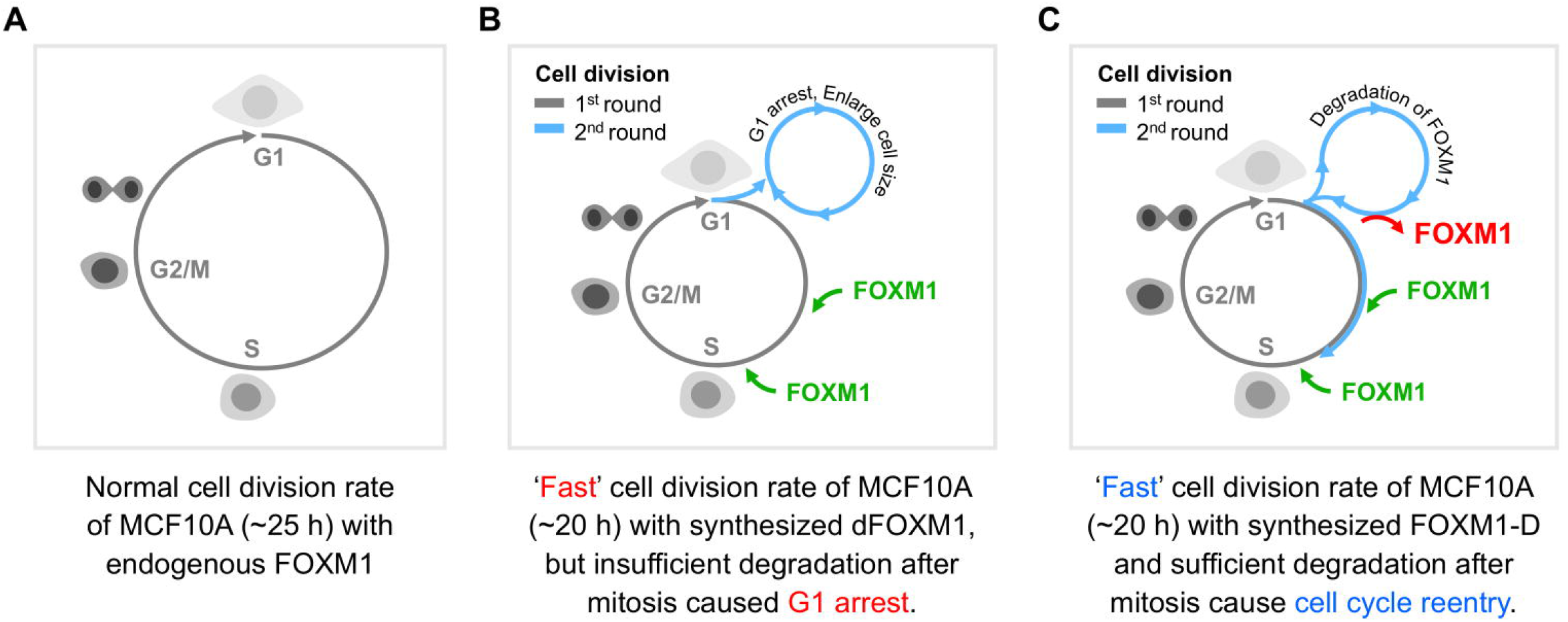
Model of FOXM1 influence on cell cycle and cell fate decision in MCF10A. **(A, B, and C)** Cell division and cell fate decision of MCF10A with the normal endogenous FOXM1 level during the cell cycle **(A)**, FOXM1 overexpressing in G1 and S phases, but insufficient degradation after mitosis **(B)**, FOXM1 overexpressing in G1 and S phases, with sufficient degradation after mitosis **(C)**.

## Materials and methods

### Cell Culture and Reagents

MCFIOA human mammary epithelial cells were obtained from ATCC (#CRL-10317, ATCC). MCF10A full growth media consisted of DMEM/F12 (#12400-024, Gibco Life Technologies) supplemented with 5% horse serum (#16050122, Gibco Life Technologies), 20 ng/Ml EGF (#AF-100-15, Peprotech), 10 μg/Ml Bovine insulin (#I1882-100MG, Sigma), 0.5 μg/Ml hydrocortisone (#H0888-1G, Sigma), 100 ng/Ml cholera toxin (#C8052-2MG, Sigma), 10,000 U/Ml Penicillin-Streptomycin (#15140122, Gibco Life Technologies). Cells were grown in an incubator with controlled temperature (37°C), CO_2_ (5%) and relative humidity (85%). To inhibit protein degradation in Figure 3D, we incubated cells with 250 μg/Ml cycloheximide (CHX, #01810, Sigma). To inhibit protein production in Figure 3C, we incubated cells with 1 μM of MG132 (#S2619, Selleck Chemicals).

### Cell Line Construction

To establish the FOXM1-DHFR-mVenus (FOXM1-D) vector, the full length of *FOXM1B* was amplified from *FOXM1* (NM_202003) Human cDNA Clone (#SC128214, Origene) then fused with the pBMN DHFR(DD)-mVenus, retroviral backbone (#29325, Addgene) (Iwamoto et al., 2010). The FOXM1^WT^-mVenus vector was constructed by fusion of the *FOXM1B* with mVenus, then cloned into PiggyBac backbone with the Histone 2B tagged with mCherry, generated by Gibson assembly (#E2611, NEB). Correct insertions for all plasmids were confirmed by sequencing.

MCF10A cells stably expressing biosensors were generated by retroviral transduction or transfection with the PiggyBac transposase system (Yusa et al., 2011). After transfection or transduction, FOXM1-D MCF10A cells were synchronized in G2/M phases under 125 nM nocodazole for 14 h, while the FOXM1^WT^-mVenus MCF10A cells were cultivated for two weeks, then sorted on BD FACSAria II to collect pure populations expressing the desired fluorescent reporters.

### Transient Perturbation of FOXM1

To transiently overexpress FOXM1 levels, we used trimethoprim (TMP)-mediated stabilization at final concentrations of 0.4, 4, 40, and 400 μM (#sc-203302, Santa Cruz). The FOXM1-D kinetic and phenotypic changes were observed after the TMP was stabilized for 12-48 h. To measure the effect of TMP withdrawal, we washed cells with fresh media 3 times to remove TMP after the desired incubation period (8 h in Figure 1E, 19 h in Figure 4J) and subsequently cultured the cells in complete media without TMP.

### Cell Cycle Synchronization

Prior to treatment, the FOXM1-D MCF10A cells were seeded at 5,000 cells/well (for G1 phase serum-deprivation), 2,500 cells/well (for S phase double-thymidine block), and 4,000 cells/well (for G2/M phase nocodazole treatment) with a complete media overnight onto poly-D-lysine-coated CellCarrier-Ultra-96 well Black (Perkin Elmer).

To synchronize in the G1 phase, we washed cells gently with serum-free media 3 times and incubated them with serum-free MCF10A media for 24 h. To synchronize in the S phase using double thymidine blockade, we first incubated cells in 2 mM thymidine (#T9250, Sigma) for 14 h. Then thymidine was removed, and cells were washed with full growth media 2 times and released into the cell cycle using complete media for 10 h. Finally, the thymidine second block was administered by incubating with 2 mM thymidine for an additional 14 h to synchronize cells in the S phase. To synchronize in the G2/M phase, we incubated cells with 125 nM nocodazole (#M1404, Sigma) in MCF10A complete media for 12-14 h. After synchronization, cells were released into the cell cycle by washing 3 times with fresh media and incubated with complete media for 12-48 h for the subsequent experiments in figures 3–4.

### EdU Incorporation and Immunofluorescence

A pulse-chase EdU assay was used to quantify cell cycle distribution and trajectory. First, cells were treated with 10 μM EdU for 15 min, and then Hoechst 33342 (Invitrogen) was added to the final concentration of 500 ng/Ml for 45 min in a CO_2_ incubator. Then, cells were fixed in 4% paraformaldehyde (PFA) for 15 min and washed with 0.1%PBS-T 3 times. Fluorescent reporter mVenus and Hoechst were imaged and collected as cycle 1 by Operetta CLS High-Content Analysis System (CLS, Perkin Elmer) using appropriate filters. Cells were then permeabilized with cold methanol for 15 min and washed with 0.1%PBS-T 3 times. The fluorescent signals from cycle 1 were bleached by fluorophore inactivation solution for 2 h or until the signals were removed relative to the background according to the CyCIF protocol (Lin et al., 2016).

After permeabilization, EdU staining was performed first to avoid bleaching other fluorescence dyes. EdU staining was performed by freshly preparing the mixture of 0.1 M Tris buffer pH 8.5 with 10 μM of carboxyrhodamine-110 azide dye, 4 mM of CuSO_4_, and 50 mM Ascorbic acid, and incubated on the sample for 1 h in the dark chamber before washed with 0.1%PBS-T, 3 times.

Only the endogenous FOXM1-labelling samples were then blocked with Odyssey PBS blocking buffer for 1 h at room temperature before incubating with the primary antibody. Rabbit-anti FOXM1 (#D12D5, Cell Signaling) was mixed with Odyssey PBS blocking buffer (LI-COR) at a ratio of 1:200 and incubated to the sample at 4°C for 24 h. After washing with 0.1%PBS-T, 3 times, samples were incubated with a secondary antibody conjugated to Alexa Fluor-488 (Life Technologies) in Odyssey PBS blocking buffer at a ratio of 1:1000 for 2 h and then washed with 0.1%PBS-T 3 times.

Finally, samples were incubated with 1 μg/Ml DAPI (#D1306, Invitrogen) for 1 h, then washed with PBS 3 times and kept in PBS at 4°C before imaging in cycle 2 (DAPI, EdU; carboxyrhodamine-110, and Alexa Fluor-488; endogenous FOXM1) by Operetta CLS.

### Live-Cell Microscopy

Cells were seeded at 4,000 cells/well onto poly-D-lysine-coated CellCarrier-Ultra-96 well Black (Perkin Elmer). Phenol red-free DMEM/F12 (#11039-021, Gibco Life Technologies) was used for serum-deprived and full-growth media preparation. In the FOXM1-D live-cell experiment, 200 ng/Ml Hoechst 33342 was added to the cells 2 h before the time-lapse experiment and added on top every 24 h to maintain the nuclear labeling signal throughout the entire duration of imaging. We captured 9 adjacent fields of images (3-by-3) with a 10% overlapping area every 10 min on a Nikon Eclipse Ti2 inverted microscope with a 20X objective, equipped with an environmental chamber controlling temperature, atmosphere (5% CO_2_), and humidity. One blank-well position (media added) was imaged every time point to enable the flat-field correction post-processing.

### Image Processing for FOXM1 quantification

In the live-cell imaging process, raw images were segmented using the open access CellProfiler Software version 4.1.2 (www.cellprofiler.org). Workflows were established for the Hoechst and mVenus channels by adding preprogrammed algorithmic modules in a pipeline. The CellProfiler pipeline can be downloaded from https://github.com/sisyspharm/FOXM1-project. There are two CellProfiler pipelines generated. First, in the correction illumination pipeline, the images from the BLANK well were used for flat-field correction using the gaussian method and smooth. The correction illumination value was collected and then used in the second pipeline for illumination subtraction.

In the second pipeline, we performed the background subtraction to the raw images by defining the average intensity of the background area outside the cell area in each frame. Then, the nuclear object was segmented by Hoechst signals and shrank for 2 pixels to identify it as the ShrinkNuc region. The cytoplasmic ring region (CytoRing) was defined by a 5 pixels-wide ring surrounding segmented nuclei (Figure 1B). The cell region was defined as a combination of nucleus and cyto-ring regions. Numerous parametric measurements were collected parallelly from the identified objects, including the total number of nuclei-stained cells, the intensity of mVenus in the various area (e.g., nuclei, cytoplasm, and cell), and cell morphological phenotypes (e.g., cell area, and cell shape). All data are represented by the mean ± SD of multiple independent experiments and paired student’s *t*-test was used to calculate the p-value.

### Image Processing for Cell Cycle Determination

The EdU-labeling immunofluorescence dataset contained two image sets; cycle 1 (Hoechst and mVenus) and cycle 2 (DAPI, EdU; carboxyrhodamine-110, and Alexa Fluor-488; endogenous FOXM1). These two image sets were first aligned by nuclear-labeling signals (Hoechst and DAPI channels), and then the object segmentation and measurements were performed. Parameters for cell cycle determination, including Nuclei area, DNA content index as DAPI intensity, and DNA proliferating index as EdU intensity, were extracted and exported in the CSV file format for both single-cell and well-average data.

### Cell Cycle Distribution

We classified the cell cycle into 4 segments (Figure 2A): G1, Early S (ES), Late S (LS), and G2/M using single-cell data described earlier. Ten thousand individual cells in asynchronous wells, each plate was first plotted as a U-shape scatter between DNA content (integrated Hoechst intensity) and DNA proliferating marker (integrated EdU intensity) for adjusting the cell cycle gate thresholding before applying it to the treatment wells. The cell cycle gate thresholding was generated by the cut-off between negative and positive staining of Hoechst and EdU using bimodal distribution in each axis. These two cut-offs clustered the cell cycle into 4 segments. The G1 phase is represented by low Hoechst and low EDU intensity. The early S phase is described by low Hoechst and high EDU, while the late S phase is characterized by high Hoechst and high EDU staining. Finally, the G2/M phase is represented by high Hoechst with low EDU cells. The process and output were performed and visualized in MATLAB (MathWorks).

### FOXM1 Trajectories in Cell Cycle

We applied the wanderlust method, and Cycler computation package (Gut et al., 2015), to the U-shape cell cycle plot for generating the cell cycle trajectories. The code is available at https://github.com/sisyspharm/FOXM1-project. To generate FOXM1 trajectories in the cell cycle, we overlaid cell cycle trajectories with the FOXM1 intensity in individual cells. The average intensity for each projectile point was calculated for further analysis.

### Mathematical Modelling

We applied a two-compartment PK-PD model to simulate protein degradation and translocation of FOXM1-D protein in G1 and G2 cell cycle phases. In the G2/M-synchronized condition, the mitotic cells were filtered out by image processing, and only attached G2-like cells were included in this study. This model consists of a simplified concept of how the FOXM1-D sensor proteins produce, translocate, and degrade. The partial differential equations for the proteins in the cytoplasm and nucleus have the following forms:

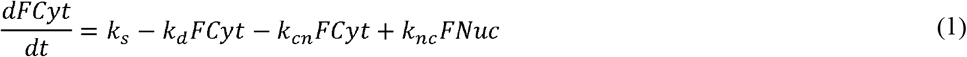

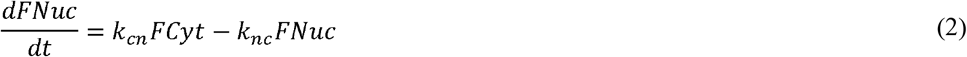

where *FCyt* is the FOXM1-D protein in the cytoplasm, and *FNuc* is in the nucleus. *k_s_* is the protein synthesis rate of the FOXM1-D protein. We assume that the proteins are synthesized only in the cytoplasm (Dahlberg and Lund, 2004) and imported into the nucleus when TMP is added (Figure 3A). *k_d_* is the protein degradation rate in the cytoplasm. *k_cn_* and *k_nc_* are the protein translocation rates from the cytoplasm to the nucleus and vice versa, respectively. Notably, this model assumes that in the absence of TMP, the levels of FOXM1-D sensors will not change over time.

### Model Estimation and Simulation

According to the mathematical model, there are four constraints of the relative rate ratio of G2 over G1 (*R_s_, R_d_, R_cn_*, and *R_nc_*) that were needed from model estimation before simulating the data. We first assumed that the rates of change between the G1 and G2 phases are unequal and that there was a linear relationship transition in the synthesis, degradation, and translocation process from the G1 to G2 phases. Firstly, we estimated the FOXM1-D synthesis rate (*k_s_*) in the cytoplasm by using the MG132, a protein degradation inhibitor, which resulted in the deletion of the degradation term in equation (3). We measured FOXM1-D in the absence of TMP, so there was no translocation process of FOXM1-D, then the translocation term was eliminated and summarized as shown in (3).

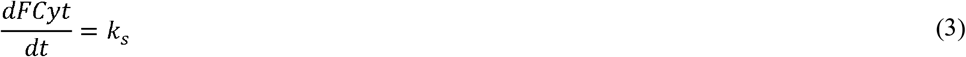

From equation (3), we fit the data after treatment with MG132 (between 4-8 h) using the linear fit package (‘poly? model in the ‘fit’ function) in MATLAB. We collected the rate ratio of G2 over G1 to represent the comparative value of *k_s_* between 2 phases (*R_s_*). Second, we estimate the degradation rate (*k_d_*) of FOXM1-D protein that occurs in the cytoplasm using the cycloheximide (CHX), the protein production inhibitor, so we eliminated the synthesis term. We hypothesized that the level of FOXM1-D from the translocation process influenced the degradation in the cytosol, so we added TMP to stabilize FOXM1-D and promoted its translocation. After 4 h of TMP stabilization, we added CHX and measured cytoplasmic FOXM1-D over time. With this strategy, we also eliminated the translocation term due to the equilibrium state of proteins, as shown in equation (4).

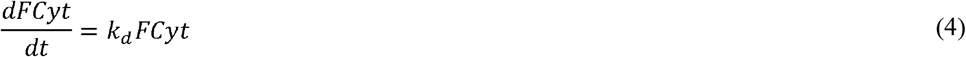

From this equation (4), we can fit the data after treatment with CHX (for 3 h) using the linear fit package (‘poly? model in the ‘fit’ function) in MATLAB. We also collected the rate ratio of G2 over G1 to represent the comparative value of *k_d_* between the 2 phases (*R_d_*). Third, to estimate the FOXM1-D translocation rate in G1 and G2 phases, we measured its dynamic during the TMP stabilization in the nuclear and cytosol area. To solve the systems of differential equations, we set up the constraints, *R_s_* and *R_d_* from previous experiments (9, 10) to solve the fitting equation (5, 6, 7, 8) as the following forms.

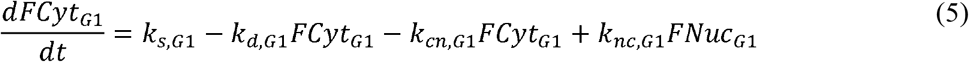

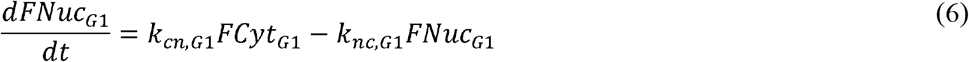

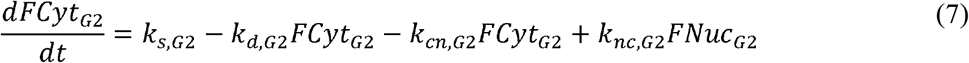

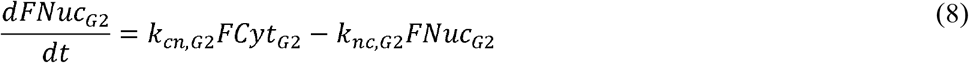

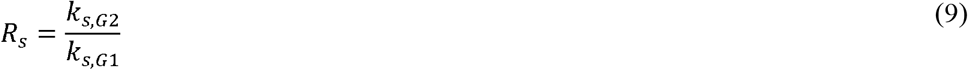

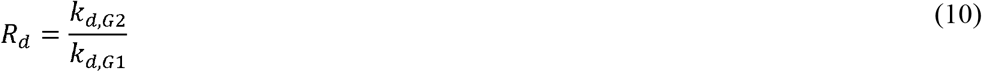

We then fitted the data in G1 and G2 synchronized conditions by restricting ±10% boundaries of constraints, using MATLAB functions ‘lsqcurvefit’ and ‘ode45’. The translocation rate ratio of G2 over G1 (*R_cn_* and *R_nc_*) was described in equations 11 and 12.

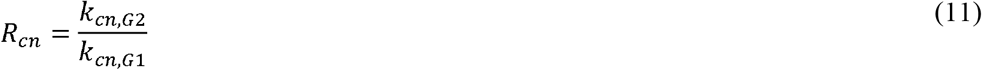

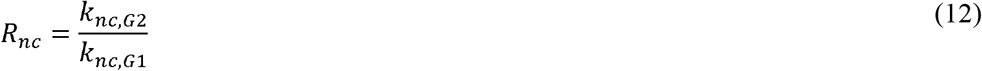

To simulate the FOXM1-D dynamic along with cell cycle progression from G1 to G2 phases, we used the single cell tracking dataset from G1 synchronized cells that progressed to G2 in the presence of TMP. We set up four constraints, *R_s_, R_d_, R_cn_*, and *R_nc_* (Table 1) prior as a limit range by accepting the ratio in ±10% boundaries. The mean intensity of nuclear and cytosol FOXM1-D in 8 single cells was used and fitted with the core model equation (1, 2). By the linear relationship between G1 and G2, we used the ‘interp? package in MATLAB and the ‘ode45’ function as solvers. The constant variables (*k_s_, k_d_, k_cn_*, and *k_nc_*) from G1 and G2 phases in the best fit were shown in table 1.

### Single-Cell Tracking and Quantification

We developed a semi-automated MATLAB-based software, ‘CellTracking’ (https://hengkp.github.com/celltracking), aimed at tracking the individual nuclei and annotating cell fate events on fast-moving cells over long timescales. In this software, we applied the ‘template matching’ and ‘whole image duplicate’ methods to automatically identify nuclei between time frames using the nearest distance with similarity of intensity and shape information. A cytokinesis event was manually identified if two nuclei in a frame wearing a similar size and intensity were linked to the same nucleus in the previous frame. Mother and daughter cells were manually linked and tracked by keeping their relationship for further analysis. The XY positions of each nucleus on the timescale were retrieved from the ‘CellTracking’ software, then aligned with the segmented objects processed by CellProfiler Software to acquire the extracted parameters as previously described. To quantify the nuclear to cytoplasmic ratio (N/C ratio) of FOXM1 activity, we used the average ShrinkNuc intensity and divided it by the average CytoRing intensity. Only cells that remained within the field of view throughout the entire duration of the experiment were considered for downstream analysis.

## Author Contributions

Conceptualization, KP, PC, and SS; Methodology, KP, PC, and MK.; Investigation, KP and PC; Formal Analysis, KP, and PC; Writing-Original Draft, KP, and PC; Writing–Review and Editing, KP, PC, EL, and SS; Visualization, KP, and PC; Supervision, SS, EL, and CP; Funding Acquisition, SS; Resources, SS. All authors contributed to the article and approved the submitted version.

## Acknowledgments

We thank Dr.Bernhard Steiert and Dr.Manupat Lohitnavy for their insightful advice regarding kinetic modeling, Dr.Siwanon Jirawatnotai for cell cycle synchronization protocols, chemical reagents, and experimental guidance, and all the members of the Siriraj Initiative in Systems Pharmacology (SiSP) lab for helpful discussions and very useful comments on this study.

## Data Availability

The data supporting the conclusion of this article will be made available by the authors, without undue reservation. Analyzed data and code is available on GitHub at: https://github.com/sisyspharm/FOXM1-project.

